# A Rat Model of a Focal Mosaic Expression of PCDH19 Replicates Human Brain Developmental Abnormalities and Behaviors

**DOI:** 10.1101/2020.06.12.145508

**Authors:** Andrzej W Cwetsch, Roberto Narducci, Maria Bolla, Bruno Pinto, Laura Perlini, Silvia Bassani, Maria Passafaro, Laura Cancedda

## Abstract

PCDH19 gene-related epilepsy or epileptic encephalopathy, early infantile, 9 (EIEE9) is an infantile onset epilepsy syndrome characterized by psychiatric (including autistic) sensory and cognitive impairment of varying degrees. EIEE9 is caused by X-linked PCDH19 protein loss of function. Due to random X-chromosome inactivation, EIEE9-affected females present a mosaic population of healthy and Pcdh19-mutant cells. Unfortunately, no mouse models recapitulate to date both the brain histological and behavioural deficits present in people with EIEE9. Thus, the search for a proper understanding of the disease, and possible future treatment is hampered. By inducing a focal mosaicism of Pcdh19 expression using *in utero* electroporation in rat, we found here that Pcdh19 signaling in specific brain areas is implicated in neuronal migration, as well as in core behaviors related to autism and cognitive function.

## INTRODUCTION

In the early development of the central nervous system (CNS), cell adhesion molecules (CAM) provide an adhesion code for cortical and hippocampal lamination (Bukalo *et al*., 2004), neural circuit wiring (Yagi and Takeichi, 2000; Gumbiner, 2005; Togashi *et al*., 2009), synapse formation (Osterhout *et al*., 2011) and maintenance of the cell-cell interactions (Nishimura and Takeichi, 2009; Harris and Tepass, 2010). Among CAMs, protocadherins (PCDHs) are characterized by spatiotemporally controlled patterns of expression for the most part restricted to specific neuronal circuits, brain nuclei, and cortical/hippocampal regions and layers (Vanhalst *et al*., 2005; Kim *et al*., 2011). This is consistent with the fact that they play specific roles in the patterning and wiring of the diverse brain areas during development. For example, restricted mutations using conditional mouse mutant lines of some of the PCDHs have revealed key roles in neuronal migration, dendritic/axon formation, and synapse function/stability in the cortex and hippocampus (Hayashi and Takeichi, 2015; Peek *et al*., 2017). Moreover, recent studies showed that the impairment of expression/function of non-clustered PCDHs (i.e., δ-Pcdhs: 8, 9, 18 and 19 among others) is related to brain disorders such as epilepsy, autism, mental disabilities, or attention deficit (Kahr *et al*., 2013). In particular, mutations in the protocadherin 19 (*PCDH19*) gene located on chromosome X (Xp22.1) cause a female-limited epilepsy (*PCDH19* gene-related epilepsy or epileptic encephalopathy, early infantile, 9 (EIEE9); OMIM # 300088).

EIEE9 is an infantile onset epilepsy syndrome characterized also by psychiatric (including autistic), sensory, and cognitive impairment of varying degrees (Depienne *et al*., 2011; Depienne and Leguern, 2012; van Harssel *et al*., 2013; Higurashi *et al*., 2015; Vlaskamp *et al*., 2019). EIEE9 people also present brain structural abnormalities. In particular, together with areas of normal architecture, focal dysplasia, heterotopia and abnormal morphology of individual neurons in the cortex as well as hippocampal sclerosis have been reported (Ryan et al. 1997; Scheffer et al. 2008; Lotte et al. 2016; Trivisano and Specchio 2018; Kurian et al. 2018; Pederick et al. 2018). Due to random X-chromosome inactivation EIEE9-affected females are composed of a mosaic population of healthy and Pcdh19-mutant cells, whereas hemizygous male carriers are asymptomatic or show much reduced psychiatric and behavioral deficits (Scheffer et al. 2008; van Harssel et al. 2013; Terracciano et al. 2012; Niazi et al. 2019). To explain gender differences, a cellular interference model has been proposed. According to this model, random X-inactivation in females leads to tissue mosaicism in which cells expressing the wild type PCDH19 protein and cells expressing a mutant PCDH19 protein co-exist and thus scramble the cell-cell communication and integration in the neuronal circuits (Ryan *et al*., 1997; Dibbens *et al*., 2008; Depienne and LeGuern, 2012; Niazi *et al*., 2019). Thus, it is the cellular interference between two populations of cells (i.e., WT and PCDH19 mutated cells) the possible cause of brain dysfunctional development, leading to symptoms in EIEE9 people (Lindhout 2008). Accordingly, the rare cases of fully affected males that have been described, arise from somatic mutations that display mosaicism (Dibbens et al. 2008; Scheffer et al. 2008; Terracciano et al. 2016; Niazi et al. 2019).

In this light, an ideal model of choice for studies on EIEE9 appears to require a cellular interference strategy. Recently, a number of mouse animal models have been generated to identify the molecular mechanism underlying EIEE9 pathophysiology. Although these animal models are being instrumental and have started to shed light on the histopathology of the disease, they have mostly failed in fully recapitulating the EIEE9 phenotype regarding overt brain malformations together with significant behavioral deficits (Lotte *et al*., 2016; Pederick *et al*., 2016, 2018; Hayashi *et al*., 2017; Lim *et al*., 2019), which is a pre-requisite for future testing of potential therapeutic treatments. Moreover, while characterized by global mosaicism in all brain areas, the current models, although valuable, did not allow to dissect which particular affected brain area was responsible for the diverse behavioural phenotypes (Lotte *et al*., 2016; Pederick *et al*., 2016, 2018; Hayashi *et al*., 2017; Lim *et al*., 2019). Here, by using *in utero* electroporation as a technique to achieve a focal mosaic of Pcdh19 downregulated cells intermixed with wild type cells in the rat cortex or hippocampus, we mimicked the focal disorganization of the brain tissue described in EIEE9-affected people. By performing histological and behavioral studies in female and male animals, we found that Pcdh19 signaling in specific brain areas is implicated, in neuronal migration, as well as in core behaviors related to autism and cognitive function.

## RESULTS

### Targeted *in utero* electroporation of PCDH19 shRNA leads to region-specific mosaicism of Pchd19 expression

To try to mimic the mosaic expression of Pchd19 in EIEE9, we reproduced a model of focal mosaicism by using the *in utero* electroporation (IUE) technique for targeting a subpopulation of neuronal progenitors of the cortex or hippocampus in rat embryos at embryonic day (E)17.5. By this means, after neuronal differentiation and completion of the neuronal migration process to the cortex or hippocampus, we were able to obtain brain area-specific cell subpopulations made of transfected neurons, interspersed with un-transfected neurons. To quantify the density of transfected *vs* un-transfected cells in the region of interest, we performed immunostaining for the cell marker Hoechst and neuronal marker NeuN of brain slices from rat pups at postnatal day (P) 9 electroporated *in utero* with a plasmid encoding for the fluorescent reported protein Td-Tomato. When we calculated the percentage of electroporated cells for layer II/III of the somatosensory cortex (Figure 1A) and the CA1 region of the hippocampus (Figure 1B), we found that the percentage of electroporated cells after counts normalization on Hoechst^+^ cells was around 5% for the somatosensory cortex and 4% for the hippocampus (Figure 1C). This percentage raised to 8% for the somatosensory cortex and 6% for the hippocampus, when we normalized the count to NeuN^+^ cells (Figure 1D).

**Figure 1.**
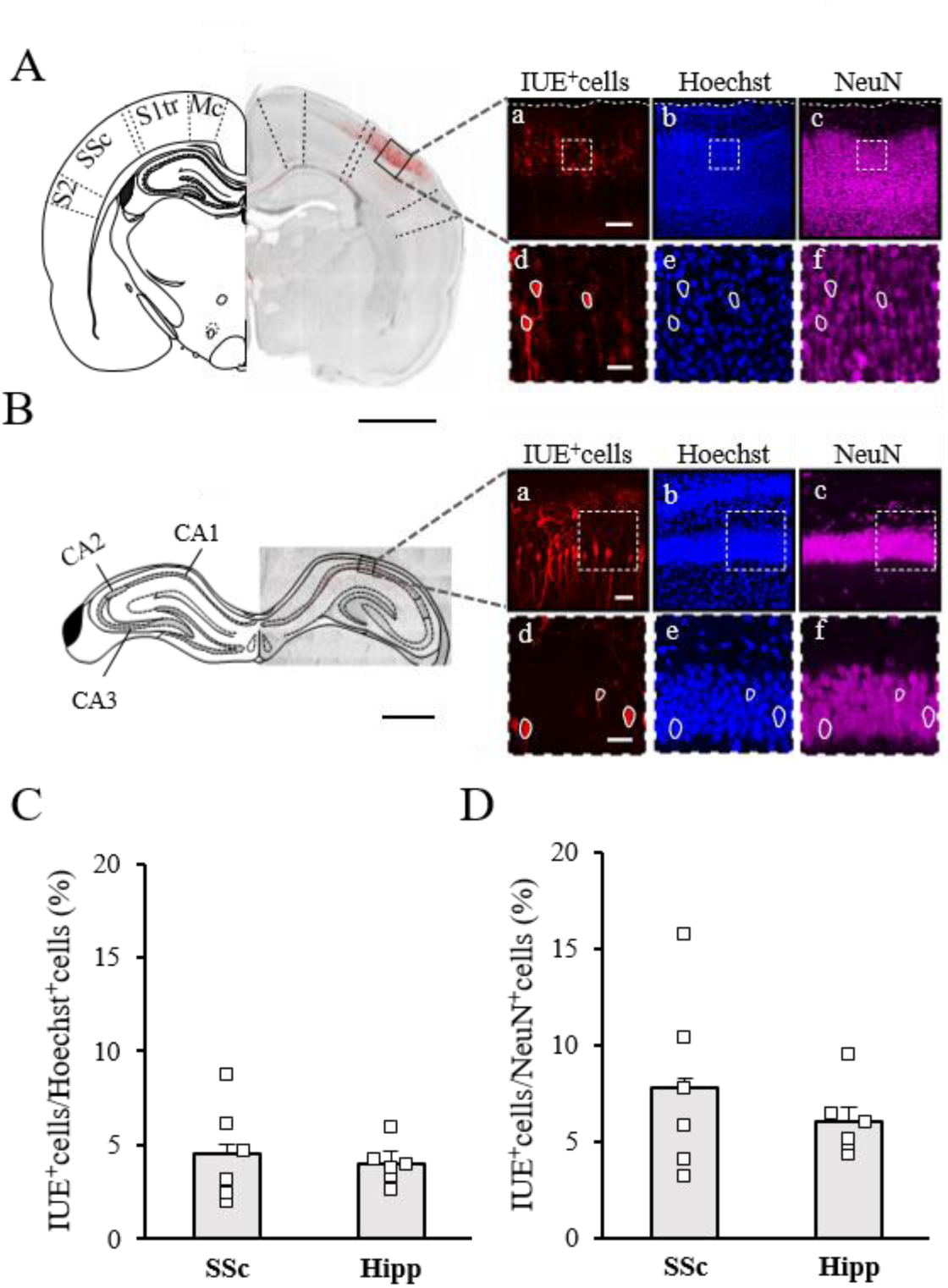
IUE of E17.5 rat embryos can specifically target or layer II/III neurons of the somatosensory cortex or CA1 neurons of the hippocampus. **(A, B)** *Left*. Example of a brain coronal slice of the somatosensory cortex (A) or hippocampus (B) from a P9 pup previously electroporated at E17.5. The slice was counterstained for Hoechst (visualized in grayscale on the left and in the blue channel in the magnification squares). Electroporated (IUE^+^) cells can be distinguished by the expression of the Tomato reporter (red channel); scale bars: 2 mm (A), 1 mm (B). The left hemisphere is replaced by a cartoon map showing the different cortical areas (A): motor cortex (MC), somatosensory cortex - trunk region - (S1tr), somatosensory cortex (SSC), secondary somatosensory cortex (S2) or hippocampal areas (B): cornu ammonis 1 (CA1), 2 (CA2) and 3 (CA3). The black square is shown magnified on the right. *Right*. IUE^+^ tomato-expressing cells (a), Hoechst^+^ cells (b) and NeuN^+^ neurons (c) are shown for the same acquisition field. Scale bars 150 µm (A), 50 µm (B). (d), (e) and (f): further magnifications of white-dashed squares shown in (a), (b) and (c) employed for calculating the IUE-density; the contour of some electroporated cells is shown in white to facilitate the overlap between the different markers. Scale bars: 30 µm (A), 30 µm (B). **(C, D)** Quantification of the average percentage of IUE^+^ cells over Hoechst^+^ (C) and NeuN^+^ cells (D) for the layer II/III of the somatosensory cortex and the CA1 region of the hippocampus, as indicated in A and B, respectively. Squares indicate values from single animals (two fields per slice and 2 slices averaged for each animal), and their averages (± SEM) are reported by bars. The images used for the different brain areas localizations were modified from “the rat brain in stereotaxic coordinates “ of Paxinos and Watson (1997).

### Pcdh19 expression is temporally and spatially regulated in the rat brain during development

We previously demonstrated that Pcdh19 shows a gradual increase of expression from E18 to P7 followed by a successive decline in the rat hippocampus (Bassani *et al*., 2018). Here, we investigated the expression of Pcdh19 during cortical development. To this aim, we performed western blot analysis on lysates of brain tissue from developing cortices at diverse time points (i.e., E15, E18, E21 and P1, P7, P16, P35; Figure 2A). Pcdh19 was poorly expressed at embryonic stages, while increased during postnatal development to reach a peak at P7 and subsequently decreased later in development and early adulthood (Figure 2B). Next, to investigate the spatial expression of Pcdh19 at its peak (P7), we performed immunostaining with a Pcdh19 antibody in coronal sections and detected Pcdh19 levels in the cortex, hippocampus and amygdala (Figure 2C). We found that Pdch19 showed a very spread pattern of expression in the motor cortex (Figure 2C (*a*)), whereas, in the somatosensory cortex, its expression was predominant in layer IV (Figure 2C (*b*)). Moreover, Pcdh19 was highly expressed in the hippocampus, especially in the stratum pyramidalis (SP; Figure 2C (*c*)).

**Figure 2.**
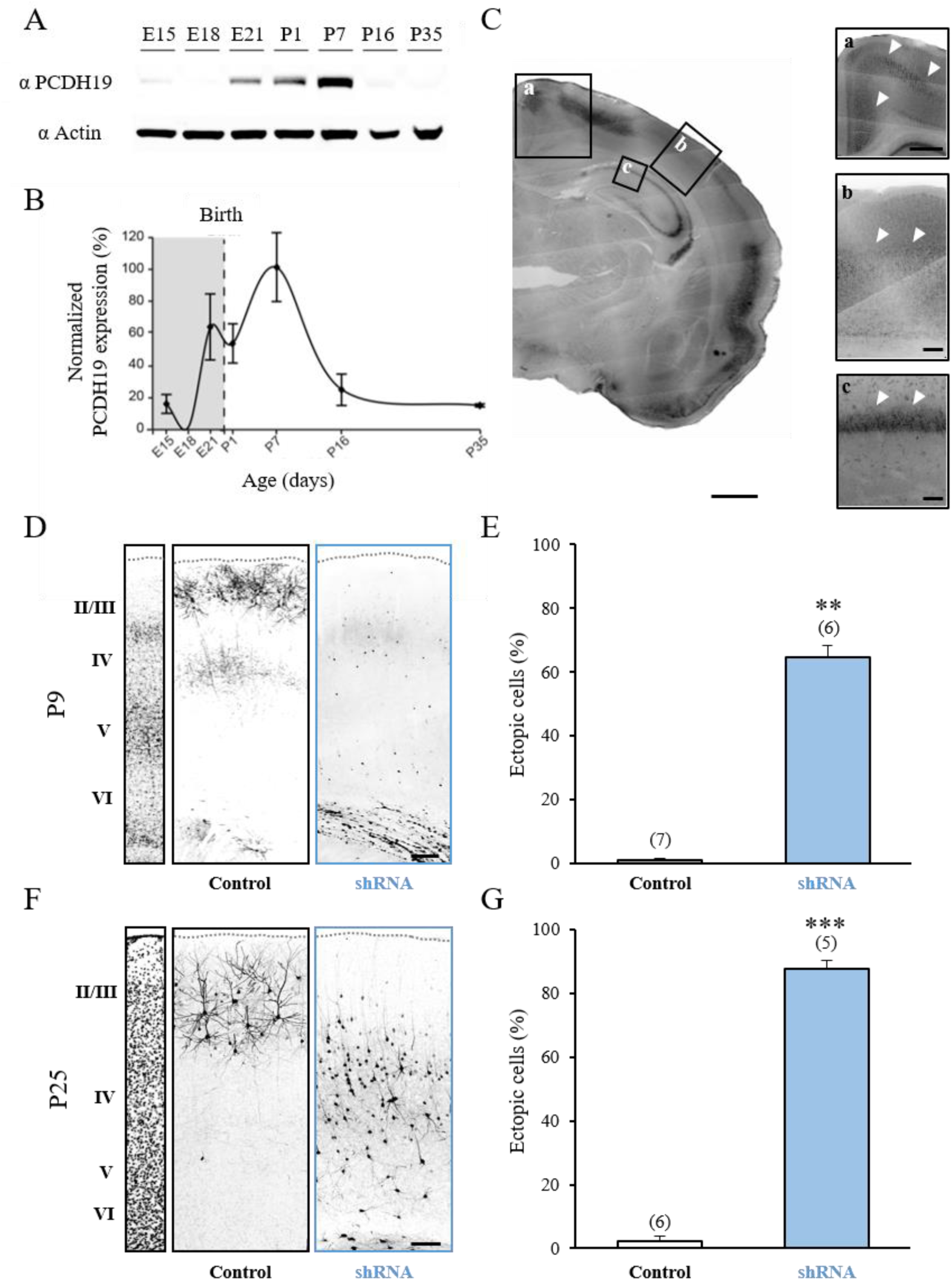
Pcdh19 is highly expressed during early postnatal development in the rat cortex and its downregulation impairs neuronal migration *in vivo*. **(A)** Representative western blot showing the temporal expression of PCDH19 in comparison to actin in lysates of rat brain cortices at different ages. **(B)** Quantification of Pcdh19 average expression (± SEM) in experiments as in A. PCDH19 signal was normalized to actin levels for each age. Data are shown as % of peak normalized PCDH19 at P7. N = 3 animals for each age. **(C)** Anti-Pcdh19 staining of a coronal section of a rat brain at P7 showing localization of Pcdh19-positive cells in the motor, somatosensory cortex, and hippocampus stratum pyramidalis (SP). Scale bar, 200 µm. (*a*) Higher magnification image of the motor cortex (as in the area indicated with squared frame) in C. White arrows point to high expression of Pcdh19. Scale bar, 50 µm. (*b*) Higher magnification image of the somatosensory cortex (as in the area indicated with a squared frame) in C. White arrows point to high expression of Pcdh19 in layer IV. Scale bar, 50 µm. (*c*) Higher magnification image of the CA1 hippocampal region (as in the area indicated with a squared frame) in C. White arrows point to high expression of Pcdh19 in SP. Scale bars, 50 µm. **(D)** Confocal images of GFP fluorescence in coronal sections of rat somatosensory cortex at P9 after IUE at E17.5 with control vector or functional Pcdh19 shRNA. Slices were counterstained with Hoechst for visualization of cortical layers (II/III, IV,V, VI, left). Scale bar, 50 µm. **(E)** Quantification of the number of ectopic cells transfected with either control vector or Pcdh19 shRNA against Pcdh19. Numbers are expressed as average percentage of the ectopic cells normalized to the total number of fluorescent cells in the same section (± SEM). Mann-Whitney test: **p < 0.01. Numbers in parenthesis: total number of animal processed (1 slice/animal). **(F)** Confocal images of GFP fluorescence in coronal sections of rat somatosensory cortex at P25 after IUE at E17.5 with control vector or Pcdh19 shRNA. Slices were counterstained with Hoechst for visualization of cortical layers. Scale bar, 100 µm. **(G)** Quantification of the number of ectopic cells transfected with either control vector or shRNA against Pcdh19. Numbers are expressed as average percentage of the ectopic cells normalized to the total number of fluorescent cells in the same section (± SEM). Student’s t-test: ***p < 0.001. Numbers in parenthesis: total number of animal processed (1 slice/animal).

Altogether, these results indicate that Pcdh19 is temporally and spatially expressed at diverse levels in different brain areas.

### Pcdh19 downregulation in the somatosensory cortex causes neuronal migration deficits and core/comorbid behaviors related to **ASD**

Focal cortical dysplasia (FCD), is a congenital abnormality of brain development where the neurons in an area of the brain fail to migrate properly (Kabat and Król, 2012). Interestingly, FCD features in people with EIEE9 and autism spectrum disorder (ASD) (Stoner *et al*., 2014; Kurian *et al*., 2018; Pederick *et al*., 2018; Trivisano and Specchio, 2018). Furthermore, around 32% of people with EIEE9 fulfill the criteria for ASD, including sensory alterations (Smith *et al*., 2018). Recently, we have demonstrated that downregulation in the upper layer of the somatosensory cortex of another cell-adhesion molecule (Negr1) with a similar pattern of expression as Pcdh19 and also associated to human ASD (Szczurkowska et al. 2018), resulted in migration defects and core symptoms related to ASD (along with sensory alterations) in rodents (Szczurkowska et al. 2018). Thus, we decided to investigate whether Pcdh19 downregulation by a short harpin RNA (shRNA) strategy (Bassani *et al*., 2018) in the somatosensory cortex resulted in migration defects and ASD-related behaviors. By IUE at E17.5, we expressed active Pcdh19 shRNAs (Bassani *et al*., 2018) or a control, scramble vector in a subpopulation of neural progenitors that would normally migrate to layer II/III of the somatosensory cortex (Dal Maschio et al. 2012; Szczurkowska et al. 2016; Cwetsch et al. 2018). Together with Pcdh19 shRNA or control vector, we also electroporated a plasmid encoding for the fluorescent reporter proteins enhanced GFP or Td-Tomato, for better visualization of transfected neurons. On electroporated animals, we examined coronal sections of the somatosensory cortex. We found that focal downregulation of Pcdh19 impaired neuronal migration *in vivo* at P9 (Figure 2D), as quantified by the number of migrating cells positioned in ectopic locations compared to controls (Figure 2D, E). Interestingly, the migration defect was maintained in early adulthood. Indeed, histological analysis of coronal sections from rat cortices at P25 electroporated *in utero* with Pcdh19 shRNA showed ectopic positioning (deep layers up to layer IV) of the neurons destined to layer II/III (Figure 2 F, G). Finally, since people with ASD may display an abnormal cortical laminar organization accompanied by alterations in dendritic spines (Stoner et al. 2014; Reiner and Dunaevsky 2015; Varghese et al. 2017; Dang et al. 2018; Szczurkowska et al. 2018), we investigated the number of spines in the neurons electroporated with Pcdh19 shRNA *vs* control littermates. In line with previous literature in layer V neurons (Hayashi *et al*., 2017), we found that Pcdh19 deficient layer II/III neurons showed a similar number of spines per micrometer in comparison to controls (not shown, Control: 0.70 ± 0.03 spines/µm, Pcdh19 shRNA: 0.71 ± 0.03 spines/µm, Mann-Whitney test, p = 0.7).

On the same set of experimental animals that we used for the histological studies, we also performed ultrasonic vocalization (USV) before sacrifice. USV is a commonly used behavioral test for social/communication deficits in ASD animal models (Fischer and Hammerschmidt, 2011). Pup littermates electroporated with either control vector or Pcdh19 shRNA were separated from the mother and littermates, and kept in isolation for 5 minutes at P9. During that time, we recorded USVs and then we calculated their frequency. We found that Pcdh19 shRNA-transfected pups vocalized significantly less in comparison to their control littermates (Figure 3A, B). Next, in another cohort of control and Pcdh19-shRNA electroporated pups, we analyzed huddling behavior, which is also considered as a social behavior in rodents (Alberts, 1978; Naskar *et al*., 2019). To this aim, each litter constituted by 10 pups-half of which were previously electroporated with Pchd19 shRNA and the rest with control construct-was separated from the mother and placed with all the other littermates in an empty area, where their interactions were video-recorded for 10 minutes (Naskar *et al*., 2019). To quantify huddling, we scored the number of clusters actively formed by pups (Figure 3C) and extracted four different parameters: Time Spent Together (the time that each pup spent with littermates in the whole huddling session), Time Spent Isolated (the time that each pup spent not in contact with littermates), Number of Different Clusters Visited (the number of cluster formed by a unique combination of littermates visited by each pup) and the Number of Cluster Switches (the number of times that pups switched from a cluster to another one). In agreement with reduced social behaviors as measured by ultrasonic vocalizations, pups transfected with Pcdh19 shRNA spent less time together (Figure 3D), more time isolated from other animals (Figure 3E) and visited less unique clusters in comparison to their control littermates (Figure 3F). No significant difference was observed in terms of cluster switches (not shown, Control: 1.60 ± 0.60, Pcdh19: 1.00 ± 0.00). Furthermore, we also examined whether Pcdh19 downregulation in the somatosensory cortex caused sensory alterations, as already described in people with EIEE9 (Smith *et al*., 2018) and as also described as comorbidity of ASD (Clarke, 2015). To this aim, we used a hot plate test at P14 (Silverman *et al*., 2010). We observed that pups transfected with Pcdh19 shRNA presented a significantly shorter latency to respond to an acute thermal stimulus when placed on a heated plate in comparison to controls (Figure 3G).

**Figure 3.**
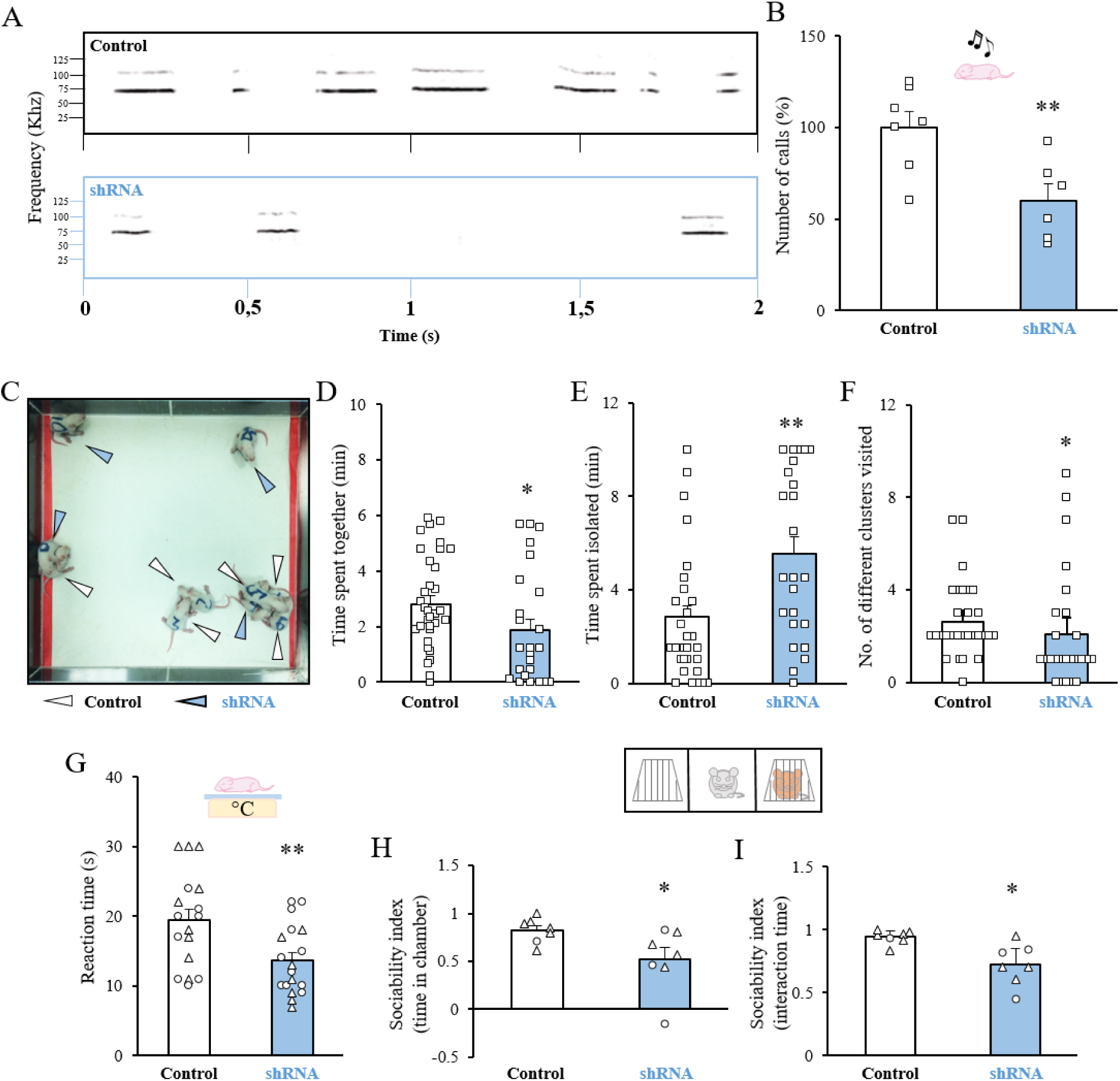
Mosaic Pcdh19 downregulation in the somatosensory cortex causes core behaviors related to autism. **(A)** Representative example of the ultrasonic vocalizations spectrograms in P9 pups *in utero* with the control vector or Pcdh19 shRNA. **(B)** Quantification of the average number (± SEM) of ultrasound vocalizations of Pcdh19 shRNA- or control vector-transfected pups normalized to values of their control littermates and expressed as a percentage in experiments as in A. Squares indicate values from single animals, and their averages (± SEM) are reported by the bars. Student’s t-test, **p <0.01. **(C)** A representative screenshot of the huddling test at minute 5, indicating P9 pups electroporated *in utero* with a control vector (white arrow) or with Pcdh19 shRNA (blue arrow). **(D)** Quantification of the Time Spent Together, expressed in minutes in huddling experiments as in C. Mann-Whitney test, *p <0.05. (**E**) Quantification of the Time Spent Isolated, expressed in minutes in huddling experiments as in C. Mann-Whitney test, **p <0.01. (**F**) Quantification of the Number of Different Cluster Visited in huddling experiments as in C. Mann-Whitney test, *p <0.05. In D, E, F, squares indicate values from single animals, and their averages (±SEM) are reported by bars. **(G)** Quantification of the latency to paw withdrawal of P14 transfected pups after placement on a hot plate. Data are presented as the average time spent (±SEM) on the hot plate until first pain reaction. Circles indicate values from single female animals and triangles indicate values from single male animal, and averages for female and male animals together (±SEM) are reported by bars. Student’s t-test, **p < 0.01. **(H)** Sociability Index based on the time spent inside the different chambers in the three-chamber sociability assay for P38-39 animals transfected *in utero* with a control vector (white arrow) or with Pcdh19 shRNA. Circles indicate values from single female animals and triangles indicate values for single male animals, and for female and male animals together (±SEM) are reported by bars. Mann-Whitney test, *p <0.05. **(I)** Sociability Index based on the time spent interacting with either a stranger mouse or an empty cage in the three-chamber sociability assay. Circles indicate values from single female animals and triangles indicate values for single male animals, and for female and male animals together (±SEM) are reported by bars. Welch ‘s t-test, *p <0.05.

Next, in analogy with our study on impaired cortical migration, we tested whether also the social behavior impairment of Pchd19 shRNA-transfected animals persisted into adulthood. In P38-39 rats electroporated *in utero* with either control vector or Pcdh19 shRNA we thus performed the three-chamber test, a classical behavioral paradigm to assess social behavior in ASD mouse models (Silverman et al, 2010; Eissa et al, 2018). In particular, we evaluated the social interactions of Pcdh19 shRNA animals or controls upon exposure to a never-met rat of the same sex (stimulus 1) *vs* an object (expressed as Sociability Index; Figure 3H, I) and to a novel rat (stimulus 2) *vs* the stimulus 1 (expressed as Social Novelty Index, not shown). We found that Pcdh19 shRNA electroporated rats spent significantly less time in the chamber where stimulus 1 was placed and also less time actively interacting (i.e. head orientation, sniffing) with the stimulus 1 in the sociability session in comparison to the control rats (Figure 3H, I;). Moreover, Pcdh19 shRNA electroporated rats showed a non-significant tendency to spend less time in the chamber where with the stimulus 2 was placed in the “Social Novelty “ assay (not shown; Social Novelty Index - time spent inside the chamber: Control: 0.22, ± 0.10, Pcdh19 shRNA: −0.04 ± 0.11; Social Novelty Index - interaction time: Control: 0.23 ± 0.13, Pcdh19 shRNA: 0.27 ± 0.14).

Altogether, these data demonstrate that Pcdh19 regulates cortical neuronal migration, and core behaviors relevant to EIEE9 and ASD.

### Pcdh19 downregulation in the hippocampus affects structural layering and impairs cognitive function

Pcdh19 is highly expressed in the hippocampus (Figure 2C (*c*)) and its downregulation impairs migration and morphology of hippocampal pyramidal neurons at P7 *in vivo* (Bassani *et al*., 2018). To investigate whether the migration defect by Pcdh19 downregulation in the hippocampus was persistent also later in life, we electroporated littermate rat embryos at E17.5 with a control vector or Pcdh19 shRNA using tripolar *in utero* electroporation (Dal Maschio *et al*., 2012; Szczurkowska *et al*., 2016). In hippocampal brain slices, we quantified the number of transfected cells located in the stratum oriens (SO) and SP at P25. While, neurons transfected with the control vector were aligned along the SP, a number of Pcdh19 shRNA-expressing neurons remained ectopically located in the SO (Figure 4A, B). Furthermore, in Pcdh19 shRNA-transfected rats, ectopic neurons were accompanied by a reduction in the thickness of the CA1 region of the hippocampus (Figure 4C, D). Moreover, hippocampal layers showed disproportions between control and treated animals (Figure 4C, E). In particular, the SO was significantly thinner (Figure 4E), whereas the stratum radiatum (SR) showed increased thickness (Figure 4E) in comparison to the control samples. Conversely, the thickness of the SP was not significantly changed (Figure 4E). Next, in analogy with our analysis in cortical cells, we performed dendritic spine counting also in hippocampal slices. In hippocampal neurons transfected with PCDH19 shRNA, we found a similar number of spine density comparison to controls (not shown, Control: 0.83 ± 0.1 spines/µm, Pcdh19 shRNA: 0.72 ± 0.05 spines/µm, Mann-Whitney test, p = 0.52).

**Figure 4.**
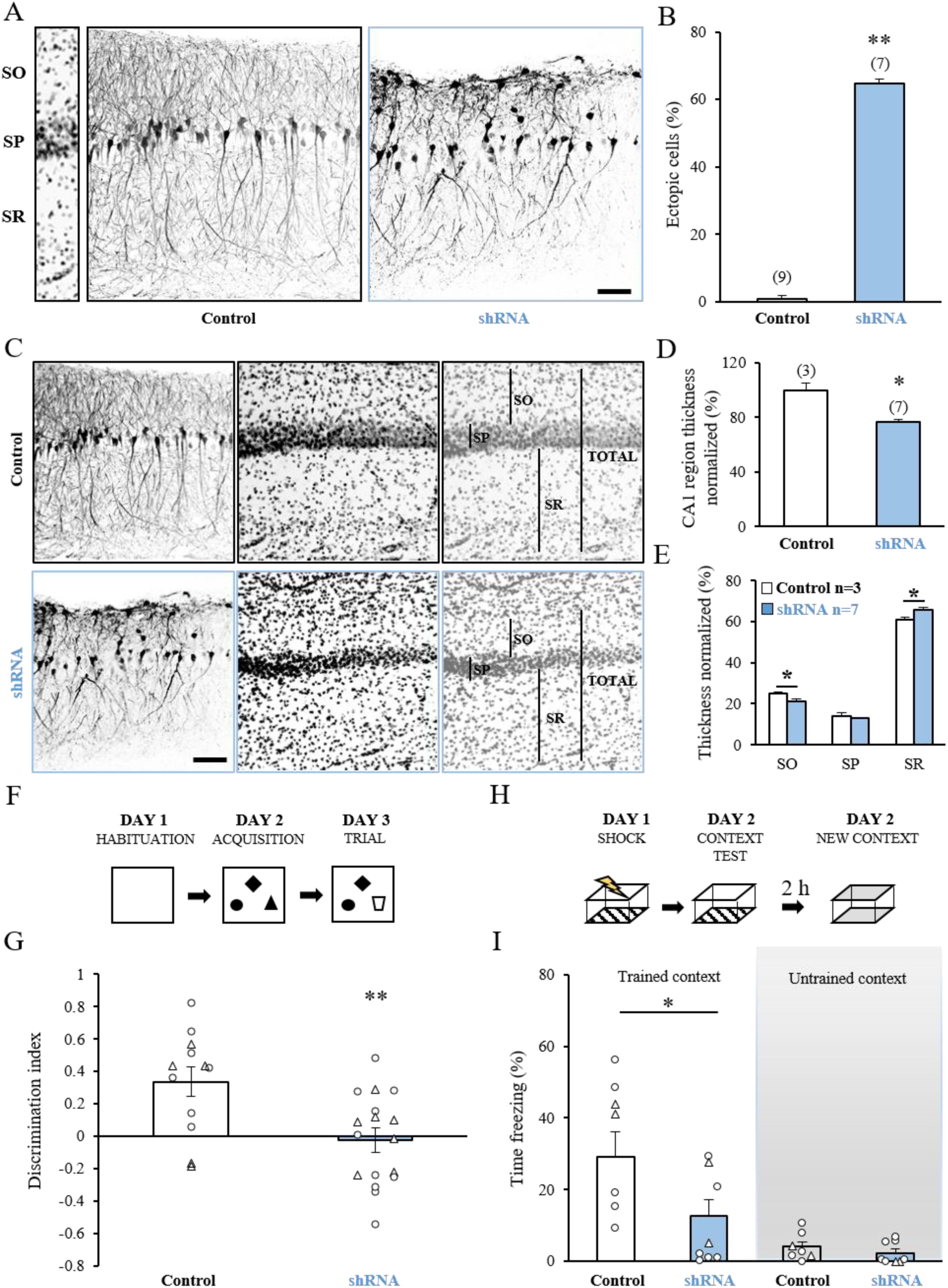
Mosaic Pcdh19 downregulation in the hippocampus impairs neuronal migration and cognitive functions. **(A)** Confocal images of GFP fluorescence in coronal sections of rat hippocampus at P25 after *in utero* transfection with control vector or functional Pcdh19 shRNA. Slices were counterstained with nuclear staining Hoechst for visualization of hippocampal layers (left). SO - stratum oriens, SP-stratum pyramidale, SR - stratum radiatum. Scale bar, 50 μm. **(B)** Quantification of the number of ectopic cells transfected with either control vector or shRNA against Pcdh19. Numbers are expressed as a percentage of the ectopic cells normalized to the total number of fluorescent cells in the same section (± SEM). Mann-Whitney test: **p <0.01. Numbers in parenthesis: total number of animal processed (1 slice/animal). **(C)** Confocal images of GFP fluorescence in coronal sections of rat hippocampus at P25 after IUE at E17.5 with control vector or Pcdh19 shRNA. Slices were counterstained with Hoechst for visualization of hippocampal layers. Black bars show thicknesses of particular layers. Scale bar, 50 μm. **(D)** Quantification of the thickness of the CA1 region in coronal slices transfected with either control vector or siRNA against Pcdh19. Numbers are expressed as a percentage normalized to the controls (± SEM); Student’s t-test: *p < 0.05). Number in parenthesis: total number of animal processed (1 slice/animal). **(E)** Quantification of the thickness of particular layers of the CA1 region in coronal slices of P25 animals transfected *in utero* with either control vector or Pcdh19 shRNA. Numbers are expressed as a percentage normalized to the total thickness of the CA1 region in each animal (1 slice/animal) (± SEM). Welch ‘s t-test: *p < 0.05. **(F)** Experimental protocol for the novel object recognition test. **(G)** Quantification of the Discrimination Index in P25 animals transfected *in utero* with control or Pcdh19 shRNA. Circles indicate values from single female animals and triangles indicate values from single male animals, and averages for female and male animals together (±SEM) are reported by bars. Student’s t-test: **p < 0.05. **(H)** Experimental protocol for the contextual fear-conditioning test. **(I)** Quantification of the freezing response in PXX animals transfected animals upon exposure to a trained (left) or untrained (right) context. Circles indicate values from single female animals and triangles indicate values from single male animals, and averages for female and male animals together (±SEM) are reported by bars. Welch ‘s t-test *p < 0.05, left; and Mann-Whitney test: p = ns, right.

Cognitive impairment ranging from mild to severe is one of the core symptoms of EIEE9 (Depienne and LeGuern, 2012; Higurashi *et al*., 2013; van Harssel *et al*., 2013; Cappelletti *et al*., 2015; Vlaskamp *et al*., 2019). Thus, we hypothesized that defective development of the hippocampus (as one of the main brain regions involved in learning and memory (Jarrard, 1993)) by Pcdh19 downregulation may contribute to cognitive impairment present in EIEE9. To test this hypothesis, we assessed control or Pcdh19 shRNA animals in two independent behavioral tests related to hippocampus-dependent cognitive functions. In particular, we first evaluated long-term explicit memory in the novel object recognition test (NOR, Figure 4F) at 4-5 weeks of age in electroporated animals. Interestingly, the animals electroporated with Pcdh19 shRNA spent significantly less time exploring a novel object in comparison to the animals electroporated with the control vector (Figure 4G), indicating poor novelty-discrimination capability. Additionally, no significant differences were observed in total object exploration or object preference (data not shown; Control: 57.48 ± 6.7 sec, Pcdh19 shRNA: 91.48 ± 8 sec; B: Trial Control: 47.44 ± 5.2 sec, Pcdh19 shRNA: 48.31 ± 6.44 sec; C: Control: object A: 39.25 ± 2.69%, object B: 30.13 ± 2.97, object C: 30.60 ± 2.73%, Pcdh19 shRNA: object A:32.62 ± 3.16%, object B: 38.88 ± 4.62%, object C: 28.49 ± 3.79%), indicating that the poor performance in the NOR was not due to alterations in total object exploration or object preference. Seven days after the last day of the NOR test, we tested the same control and Pcdh19 shRNA animals also for associative memory in the contextual fear-conditioning test (CFC; Figure 4H). The freezing behavior in response to an electric shock was scored and compared between the experimental groups. We found that animals transfected with Pcdh19 shRNA showed significantly impaired memory in the CFC test, as demonstrated by a strong reduction of the freezing response elicited upon re-exposure to the training context 24 hours after conditioning. Pcdh19 shRNA animals showed a similar freezing behavior in comparison to control animals in an untrained context (Figure 4I), indicating that both groups did not associate changed environment with the stressful stimulus.

Altogether, these results indicate that Pcdh19 downregulation in the hippocampus is associated with hippocampal structural malformation and impairment in cognitive functions relevant to EIEE9.

## DISCUSSION

Thanks to IUE as a technique to achieve a focal mosaic of WT and Pcdh19-mutated cells, we demonstrated here that Pcdh19 in the rat cortex and hippocampus is required for proper neuronal migration, core behaviors related to ASD and cognitive functions, which are altogether consistent with EIEE9 phenotype in people.

PCDH19 gene is localized on chromosome X. As a result of random X chromosome inactivation, *PCDH19* has thus a mosaic pattern of expression in females (Depienne and Leguern, 2012). In this context, we proposed here a novel approach to mimicking the mosaic pattern of expression of Pcdh19 using IUE. In particular, we used this technique to achieve focal transfection of only a certain number of progenitors in the region of electroporation and generate a mosaic of Pcdh19 and WT cells. Interestingly, while focal cortical dysplasia, ectopic neurons in the cortex, and hippocampal sclerosis were reported in EIEE9 people (Ryan *et al*., 1997; Kurian *et al*., 2018; Pederick *et al*., 2018), no major brain developmental abnormalities were observed in Pcdh19 knock out or heterozygous mice (Lotte *et al*., 2016; Pederick *et al*., 2016, 2018; Hayashi *et al*., 2017; Lim *et al*., 2019) (Table 1). Neuronal migration upon Pcdh19 downregulation has not been tested in the other EIEE9 mouse models (Lotte *et al*., 2016; Pederick *et al*., 2016, 2018; Hayashi *et al*., 2017; Lim *et al*., 2019) (Table 1). Here, in electroporated rats, we showed that downregulation of Pcdh19 in the subpopulation of neural progenitors destined to the upper layers of the developing cortex or progenitors of principal neurons of the developing hippocampus caused a significant migration delay and ectopic neurons at P9 that persisted later in adulthood. This is in line with motility studies on Pcdh19 null neurons *in vitro* (Pederick *et al*., 2016) and other studies *in vivo*, where loss of Pcdh19 expression disrupted migration in the rat hippocampal formation (Bassani *et al*., 2018) or during the neurulation in zebrafish (Biswas *et al*., 2010). The mechanisms by which PCDH19 regulates neuronal migration are still unknown. However, as for other CAMs (Kapfhammer and Schwab, 1992), it is possible that the observed migration impairment may be caused by lack, incorrect expression or simply misfunction of the extracellular domain of Pcdh19 on the surface of migrating neurons resulting in miscommunication with other cells/adhesion molecules. Interestingly, most of the mutations of *PCDH19* in EIEE9 people are related to extracellular domains responsible for the interactions with partner cells (Yang *et al*., 2019).

**Table 1.**
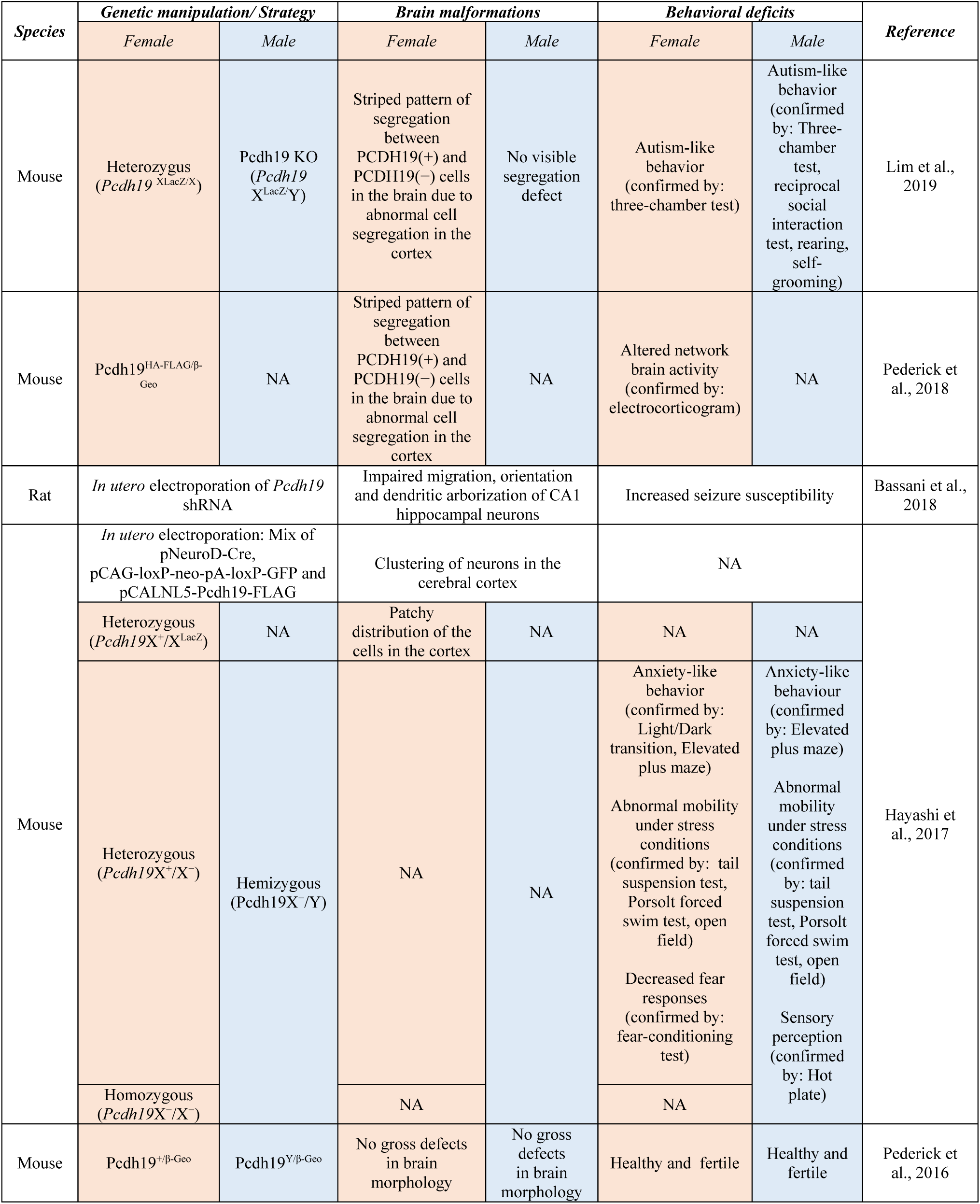
Summary of the available animal models for EIEE9 by Pcdh19 loss of expression. NA: data not available

In our EIEE9 rat model, together with the deficits in brain development, we also described behavioral phenotypes in terms of increased susceptibility to seizures (Bassani *et al*., 2018) and here social behavior, somatosensory sensation and cognitive function. Among the known rodent (i.e., mouse) models for the Pcdh19 dysregulation that reported milder brain developmental phenotypes, only some displayed also behavioral phenotypes (i.e., ASD-related behaviors in Pcdh19 X^LacZ^/X, Pcdh19X^+^/X and Pcdh19X^-^/Y mice; (Hayashi *et al*., 2017; Lim *et al*., 2019)), whereas in some others, behaviors were not investigated (i.e., Pcdh19^HA-FLAG^/^β-Geo^ and Pcdh19^+/β-Geo^; (Pederick *et al*., 2016, 2018); Table 1). Conversely, some other models, (i.e., Pcdh19 X^LacZ^/Y; (Lim *et al*., 2019)) did not show gross brain developmental abnormalities, but they did show some ASD-related behavioral phenotypes (Table 1). We found here that in rats electroporated with shRNA-Pcdh19 in the somatosensory cortex, brain developmental abnormalities were accompanied by decreased vocalisation and huddling behaviour in young pups, and poor sociability in adult animals, as already shown in animal models of ASD (Wöhr and Scattoni 2013; Scattoni, Crawley, and Ricceri 2009; Ferhat et al. 2016; Szczurkowska et al. 2018; Naskar et al. 2019). Interestingly, also a large number (32%) of people with EIEE9 carry a diagnosis of ASD (Clarke, 2015; Smith *et al*., 2018)) with social deficits early in life that persist and may become more prominent in adulthood (Depienne *et al*., 2011).

Formal diagnosis of ASD of people with EIEE9 also includes aberrant sensory perception, with both hypo- or and hypersensitivity to pain reported as a comorbidity (Clarke, 2015; Smith *et al*., 2018). Here, we found that pups with Pcdh19 downregulation in the somatosensory cortex showed faster paw withdrawal (thus, hypersensitivity) in comparison to their control littermates in the hot plate test. In particular, the effect seemed higher in male than in females (Females; Control: 16.75 ± 1.9 s, Pcdh19 shRNA: 14.81 ± 1.54 s; *Males*; Control: 21.77 ± 2.41s, Pcdh19 shRNA: 11.85 ± 1.63 s). This is in agreement with the only mouse model of EIEE9 (i.e., Pcdh19X^-^/X^+^ and Pcdh19X^-^/Y) where pain sensitivity was tested and only males showed a decreased pain tolerance (Hayashi *et al*., 2017)Table 1).

Finally, although cognitive dysfunction is one of the core symptoms of EIEE9 (Depienne and Leguern, 2012; Higurashi *et al*., 2013; van Harssel *et al*., 2013; Cappelletti *et al*., 2015; Vlaskamp *et al*., 2019), only one animal model, (i.e., Pcdh19 heterozygous females (*Pcdh19*X^+^/X^-^), (Hayashi *et al*., 2017)) showed decreased responses in the fear conditioning test (Hayashi *et al*., 2017), whereas hemizygous males (*Pcdh19*X^-^/Y) did not show alterations (Hayashi *et al*., 2017) and the other models have not been tested (Lotte *et al*., 2016; Pederick *et al*., 2016, 2018; Lim *et al*., 2019)Table 1). Here, we found that rats with Pcdh19 downregulation in the hippocampus and subjected to two independent cognitive tests showed a significant decrease in long-term cognitive functions (Betancur *et al*., 2009; Christensen *et al*., 2010; Depienne *et al*., 2011; Depienne and LeGuern, 2012; Kahr *et al*., 2013). Finally, we note that our rat model here and in (Bassani *et al*., 2018), in contrast to all the available mouse models of EIEE9, was the only able to recapitulate the main phenotype of EIEE9, which is seizure susceptibility. Whether this difference depends on how the diverse models are generated or on the animal species (rats here and in Bassani et al., 2018 *vs* mouse in the other literature), will need further investigations.

Thus, EIEE9 is a complex disease characterized by a number of behavioral symptoms, and no animal model so far has recapitulated all aspects of the pathology, but each model has proven useful to understand some specific aspect of the pathophysiology of EIEE9. Here, to the growing arsenal of mouse models of EIEE9, we add a rat model of mosaic Pcdh19 downregulation, which has been developed by using IUE and proved useful for studies on PCDH19 brain function and EIEE9. This model demonstrated to be technically easy, very versatile, quick and economical to setup (Szczurkowska et al. 2016; Dal Maschio et al. 2012; Cwetsch et al. 2018), and able to replicate the main brain morphology and behavioral feature of EIEE9. Moreover, since our model presents both developmental brain abnormalities and behavioral phenotypes related to EIEE9, it may be useful to test new therapeutic approaches aimed at rescuing the brain developmental trajectory and/or related behavioral phenotype in EIEE9. For example, the modulator of GABAA receptor activity Ganaxalone (GNX) is currently in a phase 2 clinical trial for EIEE9 (Clinical Trials Identifier: NCT03865732), but it has never been tested in animal models of EIEE9, possibly for the lack of seizure phenotype in previous mouse models. Our rat model may help in testing GNX for improvement in seizure susceptibility. Moreover, given the fundamental role that GABAA signaling exerts in brain development (Wu and Sun, 2015), ASD-related behaviors (Coghlan *et al*., 2012) - including pain sensitivity (Clarke, 2015; Smith *et al*., 2018) - and regulation of cognition (Möhler, 2009), our rat model may help in testing the ability of GNX to rescue brain morphological maturation (when given to pups), and/or accompanying EIEE9-related phenotypes.

In conclusion, we presented a new animal model for efficient and detailed studies on the mechanism underlying brain development and behavioral abnormalities due to Pcdh19 deficiency using cellular interference *in vivo*. This animal model will possibly also help to design and/or pre-clinically validate future therapeutic approaches to rescue increased susceptibility to seizure, core and comorbid behaviors related to ASD, and cognitive impairment in EIEE9 people.

## METHODS

### Animals

All animal procedures were approved by IIT licensing in compliance with the Italian Ministry of Health (D.Lgs 26/2014) and EU guidelines (Directive 2010/63/EU). A veterinarian was employed to maintain the health and comfort of the animals. Sprague Dawley (SD) rats were housed in filtered cages in a temperature-controlled room with a 12:12 hour dark/light cycle and with *ad libitum* access to water and food. All efforts were made to minimize animal suffering and use the lowest possible number of animals required to produce statistical relevant results, according to the “3Rs concept”. For animals electroporated with the experimental Pcdh19 shRNA, littermates electroporated with control plasmids were used in the same session as controls. In all experiments we used males and females.

### Western Blot

Rat cortices from were dissected in cold PBS on ice and lysed immediately in lysis buffer (2% SDS, 2 mM EDTA, 10 mM Hepes, pH 7.4, 150 mM NaCl) with 1 mM PMSF, 10 mM NaF, 1% protease and phosphatase inhibitor cocktails (Sigma). Lysates were then sonicated and clarified by centrifugation (15 minutes at 20000xg). The protein concentration of samples was estimated with BCA kit (Pierce). Equivalent amounts of protein were loaded on 10% polyacrylamide NuPAGE precast gels (Invitrogen) and subjected to electrophoresis. Next, gels were blotted onto nitrocellulose membranes (Whatman), and equal loading of proteins were verified by brief staining with 0.1% Ponceau S. Membranes were blocked for 1 hour in 5% milk in Tris-buffered saline (10 mM Tris, 150 mM NaCl, pH 8.0) plus 0.1% Tween-20 and incubated overnight at 4 °C with primary antibodies for: anti-actin (1:5000, Sigma) anti-PCDH19 (1:1000, Novus). Membranes were next washed and incubated for 1 hour at room temperature with peroxidase conjugated anti-rabbit (1:5000, BioRad), or anti-mouse (1:5000, BioRad). Stained membrains were developed with SuperSignal West Pico chemo luminescent substrate. Bands were later quantified by measuring the mean intensity signal using ImageJ.

### Generation of Constructs

shRNA#1 was a gift from Maria Passafaro laboratory. shRNA #2 was design accordingly to Elbashir, Lendeckel, and Tuschl 2001. shRNA#2; 21 nucleotide target sequence was chosen with the aid of the BLOCK-iT™ RNAi Express Sofware (Invitrogen). Both shRNA target sequences were chosen in different regions of the mRNA and their specificity for the mRNA of interest was verified by BLAST aligning with nr database. shRNAs were synthetized and cloned into pLVTH vector expressing GFP; shRNA#1: 5 ’-GAGCAGCATGACCAATACAAT -3 ’ gift from Maria Passafaro; shRNA#2: 5 ’-GCTTCTGCCCTTGTCCTAA -3 ’. As control, we used the scrambled shRNAs with following sequences: 5 ’-GCTGAGCGAAGGAGAGAT - 3 ’ and 5 ’-GCCCATCCTTCGCGTTATT -3 ’ for shRNA#1 and shRNA#2 respectively. shRNA#1 was validated in Bassani et al., 2018. shRNA#2 was co-transfected with a construct expressing rat PCDH19 in COS7 cell line and the expression of pcdh19 was assessed by western blot (data not shown: Empty vector: 82.89±9.1%; Scarmbled shRNA: 100 ±5.45%; shRNA: 9.48±2.34%). shRNA#2 significantly downregulated Pcdh19 expression (One-way ANOVA; post-hoc Holm-Sidak: **p < 0.01).

### In Utero Electroporation

All care of animals and experimental procedures were conducted in accordance with the IIT licensing and the Italian Ministry of Health. Surgeries were performed following published protocols (Dal Maschio *et al*., 2012; Szczurkowska *et al*., 2016). Briefly, E17.5 timed-pregnant Sprague Dawley rats (Harlan Italy SRL, Correzzana, Italy) were anesthetized with isoflurane (induction, 3.5%; surgery, 2.5%), and the uterine horns were exposed by laparotomy. The mix of shRNA#1 and shRNA#2 (1:1) or corresponding control scrambled shRNAs (4-6 µg/µl in water) plus pCAGGs IRES GFP (0.5 µg/µl) and the dye Fast Green (0.3 mg/ml; Sigma) was injected (5-6 µl) through the uterine wall unilaterally. In behavioral experiments, pCAG-tdTomato construct (0.5 µg/µl) substituted pCAGGs IRES GFP and was combined with the control shRNA mix for easier identification of the treated *vs* control animals. After the injection, the embryo ’s head was placed between tweezer-type circular electrodes (10 mm, somatosensory cortex electroporation) or tweezer-type circular electrodes (10 mm) and a third additional electrode (5 × 6 mm, hippocampus electroporation). For the electroporation protocol, we applied 5 electrical pulses (amplitude, 50 V; duration, 50 ms; intervals, 150 ms) by a square-wave electroporation generator (ECM 830 device; BTX Massachusetts, United States). After electroporation, the uterine horns were returned into the dam ’s abdominal cavity, and embryos allowed continuing their normal development.

### Histology and Immunostaining

Brains were fixed by transcardial perfusion of 4% PFA in PBS, cryopreserved in 30% sucrose, and then frozen and sectioned coronally (80 µm thickness) using a microtome with freezing unit (Microm HM 450 Sliding Microtome, Thermo Scientific). Free-floating slices were permeabilized and blocked with 0.3% Triton X-100 diluted in PBS and 10% NGS. Then brain slices were incubated with the primary antibodies anti-PCDH19 (Rabbit, 1:500, Novus), anti-GFP (Mouse, 1:600, AbCam or Chicken, 1:600, AbCam), or anti-NeuN (Rabbit, 1:500, Cell Signaling Technology) in 0.3% Triton X-100 diluted in PBS and 5% NGS overnight at 4°C. Immunostaining was detected using fluorescent secondary antibody anti-mouse (Alexa 488, 1:600, Thermo Fisher) or anti-rabbit (Alexa 568 and 647, 1:600, Thermo Fisher) in PBS containing 0.3% Triton X-100 and 5% NGS for 2 hours at the room temperature. Slices were counterstained with Hoechst (1:1000 Sigma). Samples were mounted in Vectashield Mounting Medium (Vector Laboratories, Burlingame, CA), and processed for fluorescent or confocal microscopy.

### Confocal Acquisition and Analysis

For the quantification of IUE-density of the somatosensory cortex (layer II/III) and hippocampus (CA1 region), images from 80-µm-thick sections counterstained with Hoechst and NeuN were acquired on a confocal laser-scanning microscope (TCS SP5; Leica Microsystems, Milan, Italy) equipped with a 63× immersion objective (numerical aperture (NA): 1.4; 2 µM thick z-stacks). To avoid excessive overlapping between neighboring Hoechst- or NeuN-positive cells with the consequent limitation of cell counting, three z-stacks (6 µm total thickness) were projected on a 2D image and cells were manually counted using the “Cell Counter “ plugin of Fiji. For each image, the electroporation density was estimated calculating the percentage of Td-Tomato cells over either Hoechst^+^ or NeuN^+^ cells. For each slice, two random fields in the central part of the electroporated region were acquired and their density values averaged. One or two slices were analyzed for each animal.

For migration analysis, images from brain sections counterstained with Hoechst were acquired on a confocal laser-scanning microscope equipped with a 10× immersion objective (NA: 0.3). Confocal images (15 µm thick z-stacks) were acquired, and Z-series were projected to two-dimensional representations. The contrast of the images was adjusted to enhance the fluorescence of cell bodies, while attenuating the signal from neuronal processes to facilitate cell counting. For the quantification of non-migrating cells, all cells in somatosensory layer II/III or hippocampal CA1 region were counted and normalized to the total number of fluorescent cells in the slice. For spine counting, confocal images were acquired using a confocal laser-scanning microscope equipped with a 63× immersion objective (NA 1.4) with 1.5× digital zoom (0.5 µM thick z-stacks) and projected on a 2D image. On each image, basal dendrites of a randomly chosen transfected neuron were visually identified, and spines were manually counted (using the “Cell Counter” plugin of Fiji) on the whole visible length of one to three secondary dendrites and divided by their respective length. Spine densities obtained for each dendrite were then averaged for each neuron. One or two images for one to three different slices were acquired per animal.

### Ultrasonic Vocalization test (USV)

All pups tested for USV were first electroporated *in utero* at E17.5 with control or Pcdh19 shRNA. Electroporated pups at P9, were separated from their mother and littermates and placed one by one in an empty container (diameter, 5 cm; height 3 cm). The empty container was then placed in a sound-attenuating styrofoam box (diameter, 30 cm; height 40 cm). 5 min recording of USVs were performed. Calls were recorded with a Microphone sensitive to frequencies of 10-180 kHz Ultrasound (Avisoft UltraSoundGate condenser microphone capsule CM16, Avisoft Bioacoustics, Berlin, Germany) and an Avisoft Recorder software. Data analysis was performed on the number of calls using Avisoft SASLab Pro.

### Huddling test

Experimental procedures were performed as previously described (Naskar et al., 2019). Briefly, each litter was formed by 10 pups, half of which were previously electroporated with Pcdh19 shRNA and the other with control vector. At P9, littermates were isolated from their mother and introduced in an empty arena (50 cm × 50 cm) where they were all separated from one another by ∼9.4 cm on the circumference of a circle (30 cm diameter) drawn on the floor of the arena. Ten-minutes videos (camera: Canon XF105 HD Camcorder, Canon; Sony HXRNC2500 AVCHD Camcorder, Sony) of freely moving pups were recorded. From the recordings, we extracted one frame every 30 seconds and visually scored the behavior of all pups in huddling groups based on their proximity and interaction. We considered a pup doing huddling when it made active and prolonged contact with one or more littermates by using his head, snout or when he formed a pile with them. A custom Python script (Naskar *et al*., 2019) was used to extract from the scores four different descriptive parameters: Time Spent Together (the time that each pup spends forming a cluster with every other littermate) Time Spent Isolated (the time that each pup spends outside of a cluster), Number of Different Clusters Visited (the number of clusters formed by a unique combination of littermates that a single pup visits during the entire huddling session) and the Number of Cluster Switches (the number of times a pup switches from one cluster to another one between two consecutive sampling intervals of 30 seconds each).

### Hot Plate test

Response to an acute thermal stimulus was measured in pups at P14 using an adapted hot plate test (Gioiosa *et al*., 2008; DeLorey *et al*., 2011). In particular, the experimenter gently placed the pup on the surface of the hot plate kept at constant temperature of 52°C. The latency to withdraw the paw from the hot plate was measured. To prevent any heat injury to pups, a cut-off latency of 30 s was applied.

### Three-chamber test

The test evaluates the social approach of the tested rat *vs* a never-met animal (stimulus 1) in comparison to an object (sociability) or *vs* a second never-met intruder animal (stimulus 2) in comparison to the stimulus 1 (social novelty). This test was performed in a similar way to that previously described (Silverman *et al*., 2010; Eissa *et al*., 2018). The three-chamber apparatus comprises a rectangle, three-chambered box of grey acrylic. The chambers are accessible by rectangle openings with sliding doors. In the first 10 minutes (habituation), the tested rat was free to explore the apparatus (Ugo Basile, Gemonio, Italy) with two empty rat cages (one in each of the two side chambers), with a cone-shaped lid to prevent the rat climbing on the top of the cages. Then, the tested rat was briefly confined in the central chamber while the stimulus 1 (previously habituated to the apparatus) was placed inside a cage placed in one of the side chambers. For the following 10 minutes (sociability test), the tested rat was allowed to explore all three chambers. Then, the tested rat was again briefly confined in the central chamber while the stimulus 2 (previously habituated to the apparatus) was placed in the other side chamber inside an empty cage. For the following 10 minutes (social novelty test), the tested rat was allowed to explore all the three chambers. The time spent exploring the object or the stimulus (interaction time) was calculated by measuring the number of seconds the mice spent showing investigative behaviour (i.e., head orientation, sniffing occurring within < 1.0 cm from the cages). In addition to the interaction time, we also calculated the time spent in the chamber, where the object or the stimulus were placed. The Sociability Index for the interaction time was calculated as the difference between the time spent investigating the stimulus 1 (T1) and the time spent interacting with the familiar object (T0) divided by the total exploration time: Sociability Index for the interaction time = (T1-T0)/(T1+T0). The Social Novelty Index for the interaction time was calculated as the difference between the time spent interacting with the stimulus 2 (T2) and the time spent interacting with the stimulus 1 (T1) divided by the total exploration time: Social Novelty Index for the interaction time = (T2-T1)/(T2+T1). Both the Sociability and Social Novelty Indexes were also calculated for the time spent inside the chambers.

### Novel Object Recognition test (NOR)

The NOR test was conducted in a gray acrylic arena (44 × 44 cm). At day 1, the rat was allowed to become habituated to the apparatus by freely exploring the arena for 15 minutes. At day 2, during the acquisition sessions, three different objects were placed into the arena and the rat was allowed to explore for 15 minutes. Object preference was evaluated during the sessions. 24 hours after the acquisition session, the rat was placed in the same arena with one object replaced by a novel one, and was allowed to explore freely for 15 minutes. We considered the exploratory behavior as direct contact with the object. In case of indirect or accidental contact with the objects, the exploratory event was not included in the scoring. The time spent exploring each object was expressed as a percentage of the total exploration time for each trial. The discrimination index was calculated as the difference between of time spent investigating the novel object (Tn) and the time spent investigating the familiar objects (Tf) over the total amount of exploration time of the novel and familiar objects: Discrimination Index = (Tn-Tf)/(Tn+Tf). The test was performed under the infrared illumination.

### Contextual Fear-Conditioning test (CFC)

Each rat was individually moved from its home cage to the fear-conditioning system (TSE Systems) consisting of a transparent acrylic conditioning chamber (44 × 44 cm) equipped with a stainless-steel grid floor. After 3 minutes, the rat received one electric shock (constant electric current; 2 s, 1.5 mA) through the floor. 15 seconds after the shock, the rat was removed from the apparatus and placed again in its home cage. On the next day, the ratwas placed in the same conditioning chamber for 3 minutes (trained context) and, 2 hours later, it was moved to a novel environment (black chamber with gray plastic floor and vanilla odor, untrained context) for 3 minutes. The freezing behavior observed in the trained and untrained context was scored and normalized on the total time spent in the chamber. The test was performed under the infrared illumination.

### Statistical Analysis

Statistical analysis was performed with Prism 7 (GraphPad Software) and R (R Core Team, version 3.6) software. All of the data are presented as the means ± SEM. Equal distribution of the variances and normal distribution of the residues were inspected by Levene ‘s and Shapiro-Wilk test respectively; if one or more violations of parametric tests assumption were detected, the corresponding non-parametric test was run. Precisely, Mann-Whitney test was performed when the normality assumption or both the assumptions were not met; when only the assumption of equality of variances was violated, Welch ’s t-test was employed instead of standard t-test.

### Data availability

All data from this work are available on the free repository OFS and can be accessed on (to be inserted)

## Acknowledgements

We acknowledge Giacomo Pruzzo, IIT Genoa for his technical support with the tripolar electrode for *in utero* electroporation. We also acknowledge Monica Morini and all the animal facility staff at IIT, Genoa, for their support.

## Author Contributions

A.W.C. and L.C. conceived, designed the study; A.W.C. and R.N. performed *in utero* electroporations and huddling, A.W.C. performed histological analysis, vocalization, novel object recognition and fear conditioning tests, M.B. and B.P. performed hot plate, social interaction, R.N. performed statistical analysis, L.P. and S.B. designed shRNA and performed Western Blots. All authors revised the manuscript. A.W.C. M.P. and L.C. wrote the manuscript;

